# Comparative transcriptomics reveals domestication-associated features of Atlantic salmon lipid metabolism

**DOI:** 10.1101/847848

**Authors:** Yang Jin, Rolf Erik Olsen, Thomas Nelson Harvey, Mari-Ann Østensen, Keshuai Li, Nina Santi, Olav Vadstein, Jon Olav Vik, Simen Rød Sandve, Yngvar Olsen

**Affiliations:** Department of Biology, NTNU Norwegian University of Science and Technology, NO-7491 Trondheim, Norway; Department of Animal and Aquacultural Sciences, Norwegian University of Life Sciences, NO-1432 Ås, Norway; BioMar AS, NO-7010 Trondheim, Norway; AquaGen AS, NO-7010 Trondheim, Norway; Department of Biotechnology and Food Science, NTNU Norwegian University of Science and Technology, NO-7491 Trondheim, Norway

**Keywords:** Wild salmon, Domestication, Vegetable oil, Circadian regulation, Long-chain polyunsaturated fatty acids, Transcriptomics

## Abstract

Domestication of *Salmo salar* has imposed strong selection for production traits since the 1970s. The domestication has also imposed a radical shift in diet. Whereas wild salmon eats invertebrates, crustaceans and fish, the dietary lipids in domestic feed has since 1990 gradually shifted from fish oil (FO) to vegetable oil (VO), causing a decrease intake of long-chain polyunsaturated fatty acids (LC-PUFA). We tested the hypothesis that this shift has induced domestication-specific features of lipid metabolism in a 96-day feeding trial of domesticated and wild salmon fed diets based on FO, VO or phospholipids (PL). We addressed this by sampling tissues central in fat uptake (pyloric caeca) and processing (liver) and quantifying RNA expression and fatty acid composition. Domesticated salmon grew faster than wild salmon, with higher gene expression in glucose and lipid metabolism pathways. The promoters of differentially expressed genes were enriched for transcription factors involved in circadian clock regulation. Domesticated salmon had lower expression of *cry2* and *nr1d1*, genes involved in negative regulation of circadian rhythm, with possible implications for the diurnal cycle of feed ingestion and basal metabolic rate. Only wild salmon showed a significant impact on growth of VO *versus* PL or FO feed, whereas domesticated but not wild salmon upregulated key LC-PUFA synthesis genes *fads2d5* and *fads2d6a* in response to VO diet. Domesticated salmon had higher LC-PUFA but lower 18:3n-3 and 18:2n-6 in liver when fed VO, suggesting that domesticated salmon can better compensate for dietary shortage of LC-PUFA compared to wild salmon.

## 1. Introduction

Domestication of Atlantic salmon started in 1971 and since then they have undergone selection in breeding programs for better growth and performance, later sex maturation, higher feed conversion rate and many other measured traits (Gjedrem, Gjøen, & Gjerde, 1991). This has led to genetic divergence, allowing domesticated salmon to grow twice as fast as wild salmon (Fleming & Einum, 1997; C. Roberge, Normandeau, Einum, Guderley, & Bernatchez, 2008). Compared to wild salmon, domesticated salmon are more aggressive during feeding in the tank environment, and have higher feed intake, feed utilization, and increased metabolic efficiency (Fleming & Einum, 1997; Christian Roberge, Einum, Guderley, & Bernatchez, 2006; Thodesen, Grisdale-Helland, Helland, & Gjerde, 1999). In addition to the targeted selection on production traits, domestication is also associated with unintentional selection on traits that are linked to the new domestic environments (Fleming & Einum, 1997; Heath, Heath, Bryden, Johnson, & Fox, 2003).

One environmental variable that changes dramatically with domestication is the feed composition and feeding regimes (Ytrestøyl, Aas, & Åsgård, 2015). In the wild, salmon is an opportunistic predator and its diet consists mostly of invertebrates in rivers, and crustaceans and small fish after they migrate to the sea. Domestic salmon on the other hand have ‘unlimited’ access to food and their diet is composed of proteins from fish and plant meal, as well as a lipid source. Up until the late 1990s this lipid source was mainly fish oil (FO) from wild fisheries. During the last two decades the FO have gradually been substituted with vegetable oils (VO) with a very different profile of long-chain polyunsaturated fatty acids (LC-PUFAs) compared to what salmon is exposed to in the wild. Lipids are not only major energy source for Atlantic salmon, but also play important roles in metabolic regulation and cell membrane function (Sargent, Tocher, & Bell, 2002). The LC-PUFAs docosahexaenoic acid (DHA, 22:6n-3), eicosapentaenoic acid (EPA, 20:5n-3) and arachidonic acid (ARA, 20:4n-6) are particularly important. They are key components of cell membranes, they regulate cell membrane fluidity, function as precursors for eicosanoid production, and are important components of neural tissues (Izquierdo, 1996; Sargent et al., 2002). It is therefore likely that modern VO based diets, with significantly reduced levels of DHA and EPA, have selected for a domestic features of lipid metabolism.

Feeding of VO-based diets naturally devoid of LC-PUFA is known to induce a compensatory response of endogenous synthesis of LC-PUFA in salmon (Datsomor et al., 2019; Stubhaug et al., 2005; Zheng, Tocher, Dickson, Bell, & Teale, 2004). Due to the past 20 years of including VO as lipid sources in salmon feed, current generations of domesticated salmon are possibly better adapted to dietary VO compared to earlier generations, but whether this involves higher LC-PUFA synthesis abilities is unclear. Another major difference between wild and domesticated diets is the levels of dietary phospholipids (PLs). PLs are important for growth and development of salmon especially for early developmental stages (Poston, 1990; Taylor et al., 2015), and dietary PL are more efficient at delivering LC-PUFA into the circulatory system and ultimately the cells compared to neutral lipids such as triacylglycerols (Cahu et al., 2009; Y. Olsen et al., 2014a). The efficiency of utilizing dietary PL could therefore also be shaped during domestication in domesticated fish, however this has never been investigated.

In the present study, we aim to explore the idea that the diet of domesticated salmon has shaped and selected for ‘domestic features’ of salmon lipid metabolism. We approach this question by running a feeding trial with both domesticated and wild salmon that are fed contrasting diets rich in either FO, VO or PL and then perform comparative transcriptomic and fatty acid analyses of two tissues involved in lipid uptake (pyloric caeca) and endogenous synthesis of LC-PUFAs (liver). Our study shows clear differences between metabolism in the wild and domestic salmon genetic backgrounds and suggests that the regulation of genes involved in circadian rhythm and lipid metabolism have been key targets during domestication related selection.

## 2. Material and methods

### 2.1 Fish, diets and experimental plan

This experiment was approved by Norwegian Food Safety Authority (Case No. 16/10070). A fast-growing strain of Atlantic salmon was kindly provided by the breeding company AquaGen AS (Trondheim, Norway). The domesticated fish have been selected for good growth and performance for 11 generations since 1971. The previous generations of domesticated salmon were always fed standard commercial diets available at the time. This means that the fish were given a freshwater diet with only marine ingredients at early developmental stages but have experienced a gradual switch in seawater diet from FO to VO since the 1990s. A group of wild salmon eggs was purchased from Haukvik Smolt AS, which is a live wild salmon gene bank situated in Trøndelag, Norway. This gene bank is operating on behalf of the Norwegian Environmental Agency to preserve genetic diversity of Norway’s wild Atlantic salmon. The parents of the wild salmon used in the present study was from the first generation of salmon originally caught in 2008 in Vosso river of Norway. Wild salmon were kept in outdoor tanks with a transparent roof, with waters that has same temperature as river. The wild fish was fed “Vitalis Røye” diet from Skretting AS (https://www.skretting.com/nb-NO/produkter/vitalis-r-ye/476027), which has an EPA + DHA content of 19-20 % of the fat, and 70 % of the ingredients are of marine origin. Approximately 1300 newly fertilized eggs of both domesticated and wild salmon were transported to hatching tanks in Ervik hatchery (Frøya, Norway). The water temperature of hatching tanks for domesticated and wild eggs were slightly different to ensure that both strains hatched and start-feed at the same time.

When the yolk sac was depleted, the wild and domestic salmon strains were separated into 12 tanks (2 fish strains x 3 diet treatments x 2 replicate tanks) with 100L water and 200 fish per tank. Feeding was initiated from the next day. The experimental tanks were randomly distributed in the hatchery and the fish of each tank were reared under same temperature, continuous light and fed 24h continuously feed every day. The fish was given three contrasting diets, either a fish oil (FO) diet high in LC-PUFA, or a plant and vegetable oil (VO) enriched diet low in LC-PUFA, or a marine phospholipid (PL) enriched diet with medium level of LC-PUFA but rich in PL (Table 1). All three diets were given to the fish from start feeding up to 94 days. To ensure sufficient DHA and EPA levels the PL used to prepare PL diet was a 50/50 mixture of krill oil (Aker BioMarine AS, Lysaker, Norway) and herring roe oil (kindly provided by Erik Løvaas from Marine BioExploitation AS, Tromsø, Norway). The feeds were produced by Sparos AS (Olhão, Portugal). The composition of the diets is shown in Supplementary Table 1. FO diets have higher DHA and ARA than PL diet, while the EPA composition was similar between the two diets (Table 1). VO diet contains higher 18:3n-3 and 18:2n-6 but lower DHA, EPA and ARA compared to the other two diets.

**Table 1.**
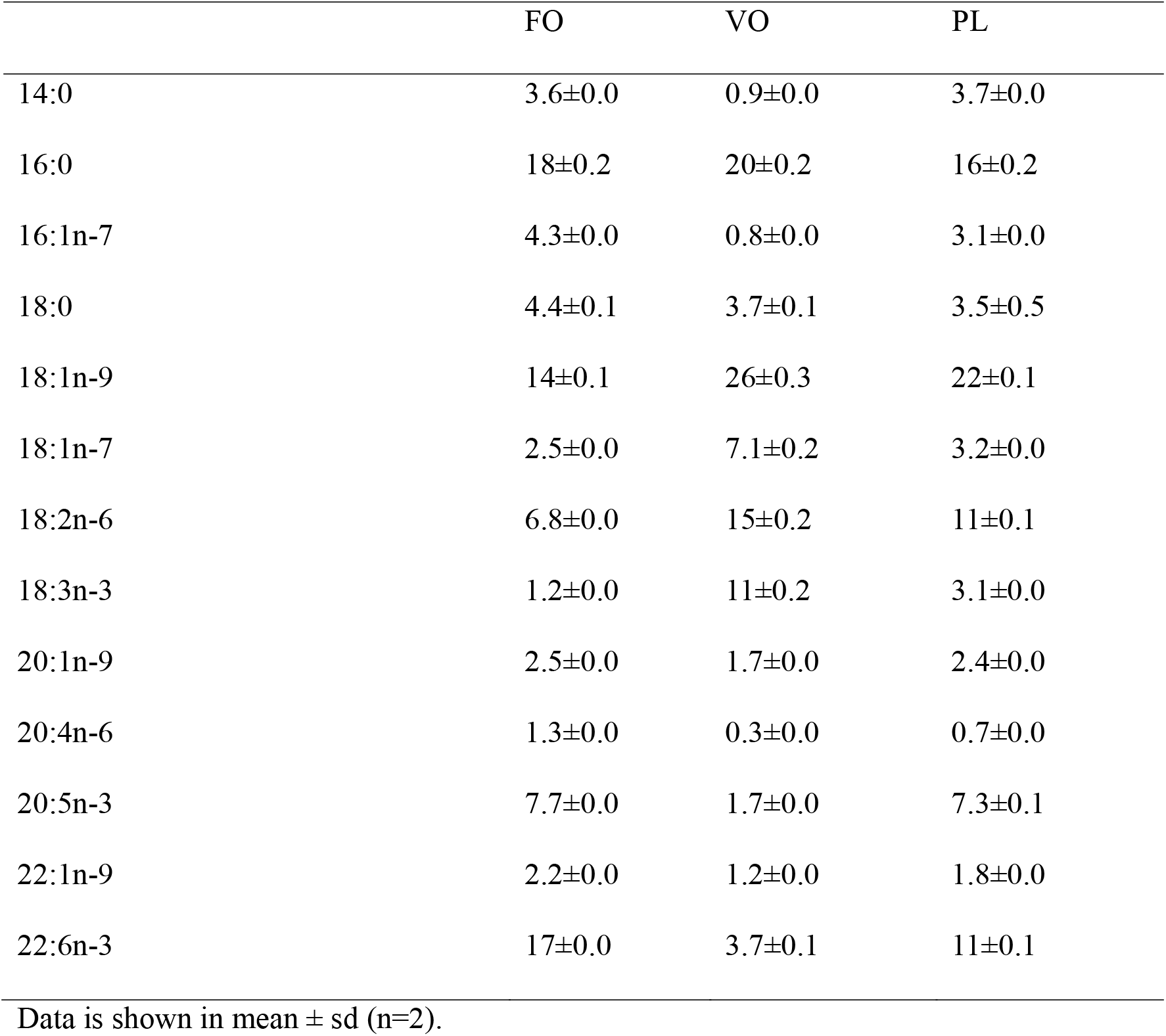
Percent of fatty acids in total fatty acids of three diets rich in fish oil (FO), vegetable and plant oil (VO), or vegetable and marine phospholipid oil (PL).

Fish was sampled after 94 days of feeding. The fish were sacrificed by exposure to 200 mg/ml Benzoak vet. (ACD Pharmaceuticals AS, Oslo, Norway), then immediately put in sterile pertri dishes and dissected under a dissecting microscope. The pyloric caeca and liver tissues were immediately transferred into 2mL Eppendorf tubes, and either filled with RNAlater and put on ice for RNA isolation, or frozen in dry ice for lipid extraction. Tissues for RNA isolation were kept at 4°C for 24h to allow sufficient penetration of the solution into the tissues, and then kept at −80 °C until RNA extraction. Tissues for lipid extraction were directly transferred to −80 °C after sampling.

### 2.2 RNA isolation and transcriptomic sequencing

Four individual fish from each tank were used for RNA isolation. The RNA extraction was performed with the RNeasy Plus Universal Kit (Qiagen, Hilden, Germany), according to the manufacturer’s instructions. The concentration and integrity of RNA were determined by a Nanodrop 8000 (Thermo Fisher Scientific, Waltham, USA) and a 2100 Bioanalyzer (Agilent Technologies, Santa Clara, USA), respectively. All RNA samples had RNA integrity (RIN) values higher than 8, which is sufficient for RNA sequencing. Sequencing libraries were prepared with a TruSeq Stranded mRNA Library Prep Kit (Illumina, San Diego, USA) according to the manufacturer’s protocol. Libraries were sequenced using 100bp single-end mRNA sequencing (RNA-seq) on Illumina Hiseq 2500 (Illumina, San Diego, CA, USA) at the Norwegian Sequencing Centre (Oslo, Norway).

The method for handling RNA-sequencing (RNA-seq) data has been described in detail in previous studies (Gillard et al., 2018; Jin et al., 2018). In brief, read sequences were quality trimmed using Cutadapt (v1.8.1) before being aligned to the salmon genome (ICSASG_v2). Raw genes counts were generated using HTSeq-counts (v0.6.1pl) and the NCBI salmon genome annotation (available for download at http://salmobase.org/Downloads/Salmo_salar-annotation.gff3).

### 2.3 Lipid class separation and fatty acid analysis

Total lipid was extracted from two individual fish from each tank by using the method of Folch, Lees, and Stanley (1957). Extracted total lipid was then applied onto 10 x 10 cm silica plates (Merck, Darmstadt, Germany) and separated by using methyl acetate/isopropanol/chloroform/methanol/0.25% KCl (25:25:25:10:9, by vol) for polar lipids and hexane/diethyl ether/glacial acetic acid (80:20:2, by vol) for neutral lipids (R. E. Olsen & Henderson, 1989). To avoid the oxidation of fatty acids, the plates were exposed to iodine vapor to visualize the lipid class for fatty acids analysis (Li & Olsen, 2017). Lipid bands of phosphatidylcholine (PtdCho), phosphatidylethanolamine (PtdEtn) and triacylglycerol (TAG) were separately scrapped out into 10mL glass tubes. Fatty acid methyl esters (FAME) of each lipid class were prepared by acid-catalyzed transesterification at 50°C for 16 hours (Christie, 1973) before quantified by a Agilent 7890B gas chromatograph with flame ionization detector (Agilent Technologies, Santa Clara, CA).

### 2.4 Data analysis

The analysis of RNA-seq data was performed in R (v3.4.1) (Team, 2013). Only genes with a minimum counts level of at least 1 count per million (CPM) in more than 25% of samples from each tissue were kept for differential expressional analysis. Differential expression was tested separately on pyloric caeca and liver using R package edgeR (Robinson, McCarthy, & Smyth, 2010). A full interaction model (Diet + Strain + Diet x Strain) was used in each tissue separately to find differential expressed genes (DEGs) between wild and domesticated salmon under any dietary treatments. Genes with a false discovery rate (FDR), an adjusted *p* value (*q*) <0.05 and absolute log2 fold change (|Log2FC|) > 1 were considered to be differentially expressed genes (DEGs) between any test conditions. KEGG ontology enrichment analysis (KOEA) was conducted using edgeR. Significant values (*p*<0.05) were generated based on hypergeometric test where the number of DEGs was compared to total genes annotated to each KO term. A test for enrichments of transcription factor binding sites (TFBS) motifs in the promoter regions (between −1000bp and 100bp from transcription starting sties) of salmon genes was done by using a hypergeometric test in the R package SalMotifDB, which is interacting with a database of transcription factor binding sites for salmonids (https://salmobase.org/apps/SalMotifDB) (Mulugeta et al., 2019).

To further investigate diet specific effect on gene expression between wild and domesticated salmon, samples of different diet were separated to be used for testing differential expression of genes between wild and domesticated salmon under each diet. Same cut-off was used (*q*<0.05 & |Log2FC| > 1) to identify DEGs. For visualize expression levels between different genes and tissues, normalized counts in the form of transcripts per million (TPM) values were generated. Raw gene counts were first divided by their mRNA length in kilobases to normalize for transcript length, and then divided by the total number of counts from each library to normalize for sequencing depth (Jin et al., 2018).

Statistical analysis of fish weight and fatty acids composition was also performed in R. Two-way ANOVA with Tukey HSD post hoc test was used to test the effect of strain and diet on fish weight and fatty acids composition of each lipid class, tissue and sampling day. Differences were considered significant when *p* < 0.05.

## 3. Results

### 3.1 Growth and development

The domesticated salmon was significantly larger than wild salmon at all sampling times (Figure 1 and Supplementary Table 2). At the end of the trial (94 days), domesticated salmon reached an average 4.5g, while wild salmon had a mean size of 2.6g. There were no significant differences in weight between domesticated salmon fed FO, VO and PL enriched diets. The growth of wild fish appeared more sensitive to different diets, with VO-fed fish being smaller than PL-fed fish at day 65 (*p* = 0.03) and day 94 (*p* = 0.02). FO-fed wild salmon showed intermediate weights, but their weights were not significantly different from either FO or PL diets. A complete table of fish weight and statistics from all time points is found in Supplementary Table 2.

**Figure 1.**
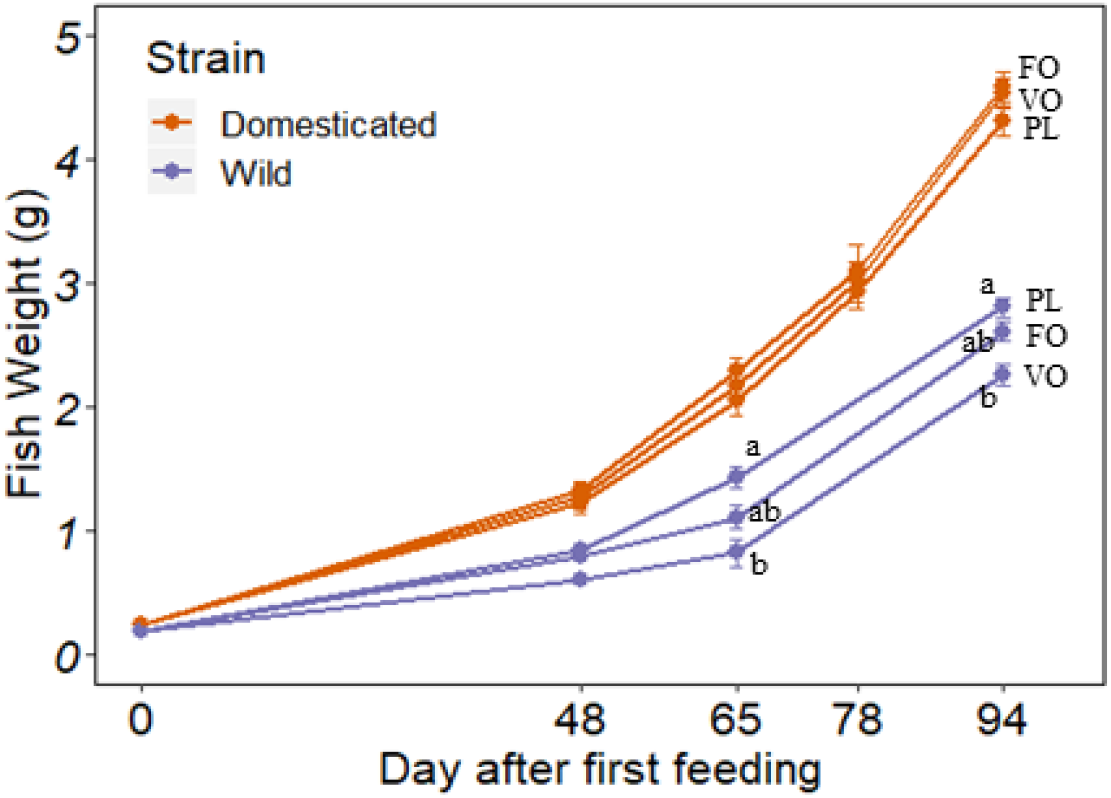
Weight of domesticated and wild salmon fed diets high in fish oil (FO), vegetable oil (VO) or phospholipid oil (PL) during early stages of development. Data are means ± SE (n>100 per group at day 93, n>20 at other days). Different letters indicate significant (*p*<0.05) different of fish weight between wild fish fed FO, VO and PL diet at day 65 and 94.

### 3.2 Transcriptomic differences between domesticated and wild salmon

On average 20 million reads were generated from each sample (min: 12M reads, max:32M reads), with ∼85% of the reads mapping to the salmon genome. Out of 81597 annotated loci, 28980 and 24119 genes passed this filtering criteria in pyloric caeca and liver, respectively. Principle component analysis (PCA) on Log2 CPM of the top 1000 most variable genes, identified a clear separation of domesticated and wild salmon in both pyloric caeca and liver (Figure 2).

**Figure 2.**
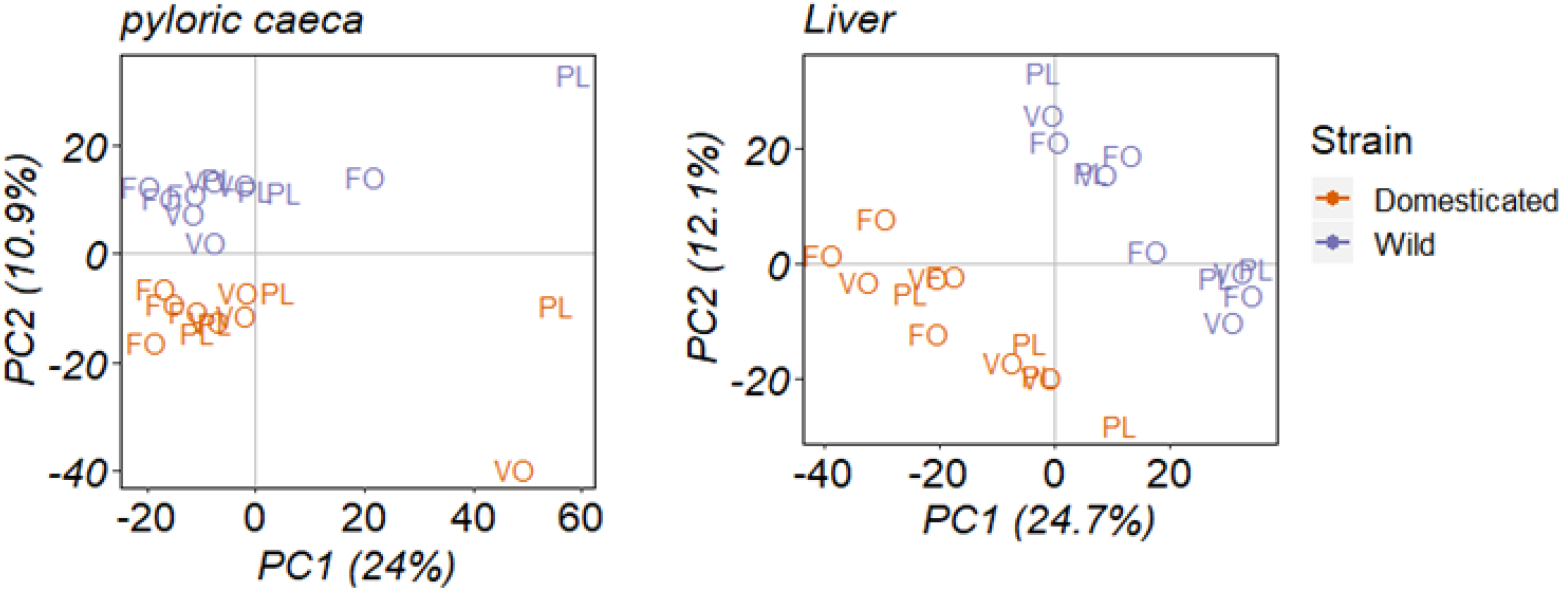
Score plot of PCA on log2 count per million (CPM) of the top 1000 most variant genes across all samples (4 replicates x 2 strains x 3 Diets). Two salmon strains (domesticated and wild) were fed either fish oil (FO), vegetable oil (VO) or phospholipid (PL) rich diets from initial feeding. Pyloric caeca and liver samples were taken after 94 days of feeding.

A full interaction model design in edgeR (Chen, McCarthy, Robinson, & Smyth, 2014) was applied to test differences in gene expression between domesticated and wild salmon across all dietary treatments on day 94. This resulted in identification of 187 differential expressed genes (DEGs) in pyloric caeca and 379 DEGs in liver between wild and domesticated salmon. The complete list of DEGs in pyloric caeca and liver is shown in Supplementary Table 3. By mapping DEGs to the KEGG database of metabolic pathways, we have identified 17 pathways that were significantly enriched (*p*<0.05) in pyloric caeca, while 11 pathways were enriched in liver (Figure 3 A and Supplementary Table 4). The DEGs in pyloric caeca were enriched in pathways for glycerophospholipid, glycosphingolipid and glycosaminoglycan metabolism, which are known to be major component of cell membrane (Figure 3 A). A number of cell-signalling pathways were also enriched, including phosphatidylinositol signalling, calcium signalling, apelin signalling, C-type lectin receptor signalling and GnRH signalling pathways. In liver, the DEGs were enriched in metabolic pathways including linoleic acid, glycolysis and gluconeogenesis, fructose and mannose, cysteine and methionine, retinol metabolism pathways (Figure 3 A). Enrich genes involved in intercellular interactions included cytokine-cytokine receptor interaction, neuroactive ligand-receptor interaction and gap junction pathways.

**Figure 3.**
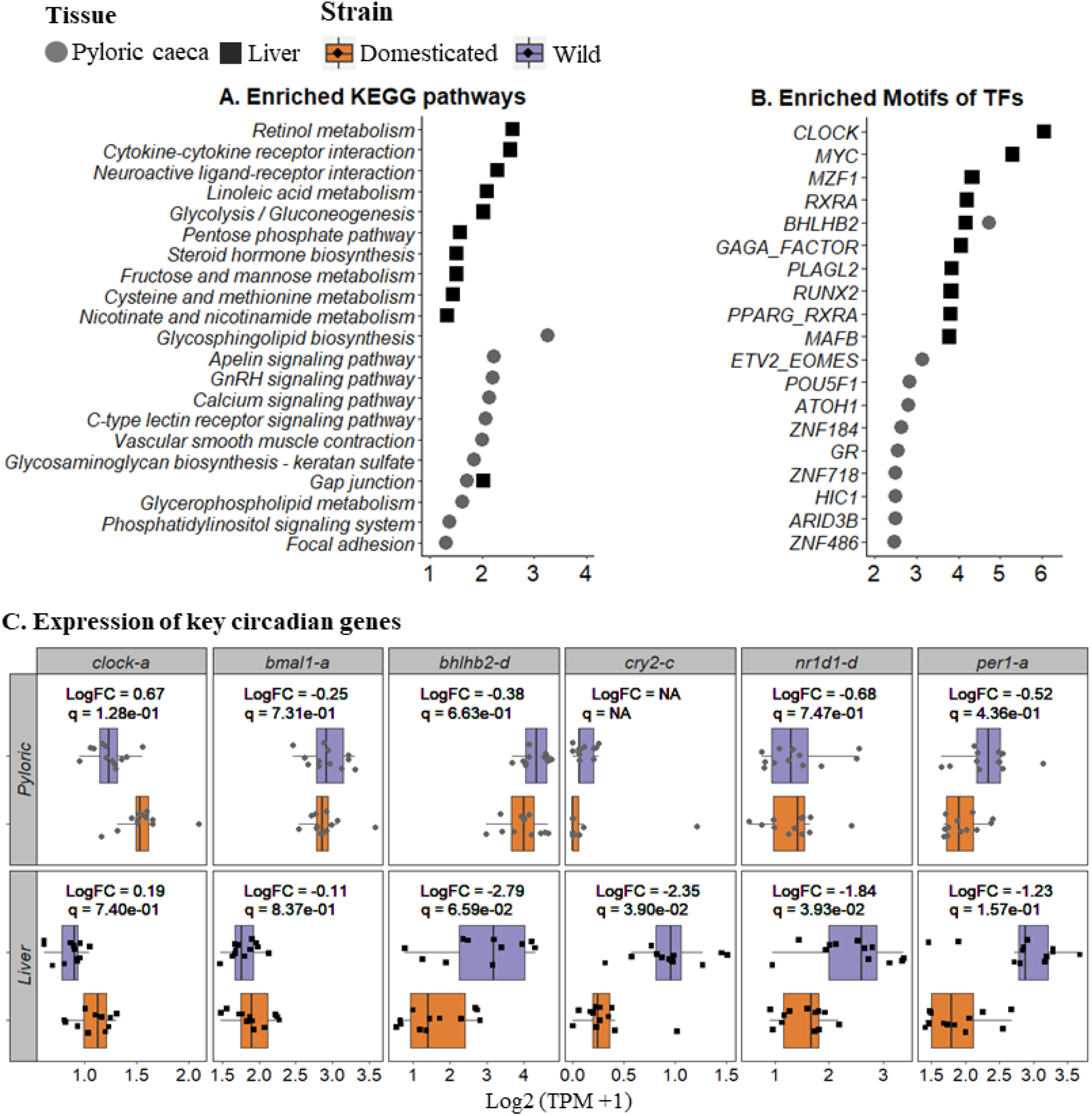
Differential expressed genes (DEGs) between domesticated and wild salmon. **A)** KEGG enrichment shows significant (*p*<0.05) enriched pathway, and proportion (%) of up/down regulated DEGs in each pathway. **B)** Motif enrichment analysis shows top 10 most significantly (*p*<0.005) enriched motifs of transcription factors in promoter regions (−1000bp to 200bp from TSS) of DEGs as compared to all expressed genes in pyloric caeca and liver. Hypergeometric test was applied on both KEGG and motif enrichment analyses, by comparing the number of DEGs to total genes annotated to each KEGG pathway or each motif. Motif enrichment analysis was done by using SalMotifDB (https://salmobase.org/apps/SalMotifDB). **C)** Expression of key circadian genes in pyloric caeca and liver of domesticated and wild salmon. Gene expression was shown in log2 transcript per million plus one (TPM +1). No statistics was shown for *cry2-c* gene in pyloric caeca, since the gene expression is too low (CPM <1) to be used for differential expression analysis.

Transcription factor binding site (TFBS) motif enrichment analysis on promoters (−1000bp to 200bp from TSS) of DEGs for each tissue resulted in 16 significant (*p*<0.005) enriched motifs in pyloric caeca and 128 enriched motifs in liver (Supplementary Table 4). The most enriched motif in pyloric caeca was BHLHB2, which is known to be involved in circadian regulation (Figure 3 B) (Dunlap, 1999). Several other enriched motifs are associated with intestinal development and cell differentiation, including ETV2 (Jedlicka & Gutierrez-Hartmann, 2008), ATOH1 (Shroyer et al., 2007), GR (Lebenthal & Lebenthal, 1999), GATA-1 (Kanki et al., 2017). The most enriched TFBS motif in liver was a CLOCK motif, which is a predicted binding motif for the master regulator of the circadian clock (Dunlap, 1999). Similar to pyloric caeca, BHLHB2 motif was also identified in the top 10 most enriched TFBS motifs in liver. In addition, three lipid metabolism related motifs (RXRA, PPARG, PLAGL2) populated the top-10 enriched TFBS list (Tontonoz, Hu, & Spiegelman, 1994; Van Dyck et al., 2007).

To further investigate differences in expression of genes linked to circadian rhythm between wild and domesticated salmon, we compared the expression of key genes encoding circadian clock related transcription factors *clock*, *nr1d1*, *bmal1*, *bhlhb2*, *per*, and *cry* (Figure 3 C and Supplementary Figure 1). A systematic difference in circadian clock gene expression was observed between livers of wild and domesticated salmon (Figure 3 C and Supplementary Figure 1), although not all genes were significant regulated at *q*<0.05. Nevertheless, the regulators (*cry2-c*, *nr1d1-a, per1-a, bhlhb2-d* and *nr1d1-a* genes) acting as suppressors of the master regulators of circadian rhythm (CLOCK/BMAL) were consistently lower expressed in domesticated salmon compare to wild (Figure 3 C). However, similar expression levels of *clock* (logFC=0.2, *q*=0.7) and *bmal1* (logFC=-0.1, *q*=0.8) genes which encodes master regulator were found between domesticated and wild salmon (Figure 3 C and Supplementary Figure 1). No difference in circadian clock gene expression was observed between pyloric caeca of wild and domesticated salmon.

### 3.4 Differential regulation of lipid metabolism genes between domesticated and wild salmon

To better understand the effect of diets on gene expression differences between domesticated and wild salmon, we compared gene expression separately between domesticated and wild under each diet. In pyloric caeca, a total number of 230 DEGs were identified between domesticated and wild salmon with FO diet, 164 DEGs were found with VO diet and 689 DEGs were found with PL diet (Supplementary Table 5). Out of these DEGs, only 8 genes were involved in lipid metabolism pathways. This includes *ptdss2* genes of phosphatidylserine synthesis, which was significantly (*q*<0.05 & |Log2FC|>1) higher expressed in domesticated salmon regardless of the dietary treatment (Figure 4). Two phosphatidylethanolamine synthesis genes, *pcyt2c-a* and *pcyt2c-b* were both higher expressed in domesticated than wild salmon when fed FO or VO diet, while no difference in gene expression was found when the fish was given PL diet. On the other hand, *etnk2-a* gene involved in phosphatidylethanolamine synthesis, was significantly higher expressed in domesticated compared to wild salmon only when the fish was given PL diet. Feeding with VO induced key genes in LC-PUFA synthesis pathway (*fads2d5* and *fads2d6a,* see Figure 4) in both domesticated and wild salmon. However, no expression difference was observed for these two genes, or any other LC-PUFA synthesis genes between domesticated and wild salmon for any dietary treatment (Figure 4 and Supplementary Table 5).

**Figure 4.**
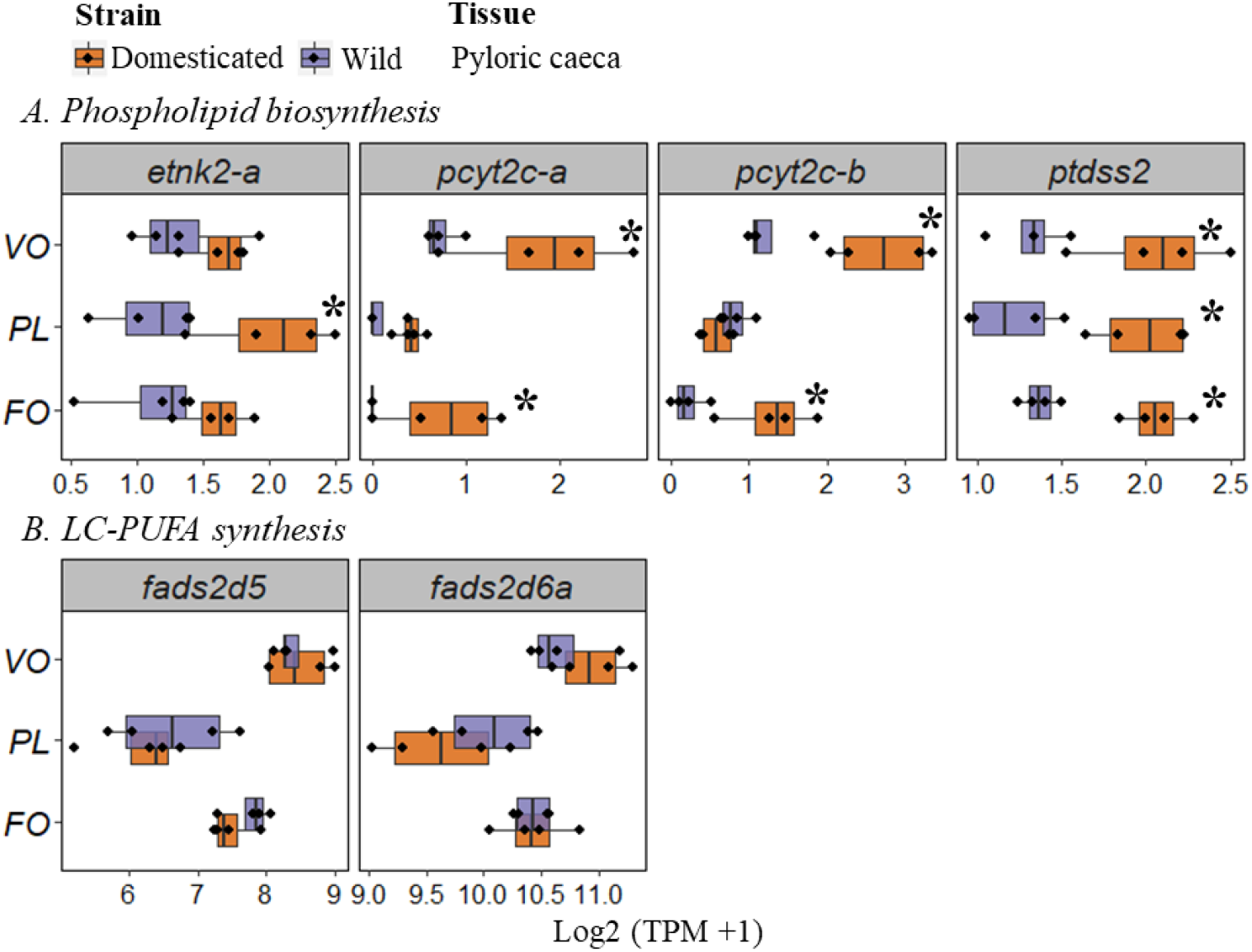
Expression of 6 genes involved in lipid metabolism in pyloric caeca of wild and domesticated salmon at day 94 after feeding either fish oil (FO), vegetable oil (VO) or phospholipid (PL) diets. Gene expression was shown as Log2 transcript per million plus one (TPM + 1) which was normalized by library size and mRNA length. Asterisk indicates differential expressed genes (DEGs, *q*<0.05 & |log2FC| >1) between domesticated and wild salmon under each dietary treatment.

The number of DEGs between liver of domesticated and wild salmon under each diet was 591 (FO), 179 (VO) and 243 (PL) (Supplementary Table 5). Liver had more DEGs involved in lipid metabolism (28) compared to pyloric caeca (8). Four DEGs in liver had significantly (*q*<0.05 & |Log2FC|>1) higher expression in domesticated compared to wild salmon under VO diet, but not under FO or PL diet (Figure 5 B). This includes genes with key functions in LC-PUFA synthesis (*fads2d5*, Log2FC = 1 & *q* = 0.02), gene involved in acyl-CoA synthesis (*acsbg2b-b*, Log2FC = 1.4 & *q* = 0.03), and fatty acid transport (*fabp7b*, Log2FC = 3.6 & *q* = 0.02). Although not significant, domesticated salmon fed VO diet also had higher expression of *fads2d6a* (Log2FC = 0.7 & *q* = 0.2) and *srebp1d* (Log2FC = 0.8 & *q* = 0.2) compared to wild salmon fed same diet, while the expression difference of the two genes was negligible when the fish was under FO or PL diet (Figure 5 A & B). A key gene involved in conversion of lipids to energy, *cpt1aa* was lower expressed (Log2FC = −1.2 & *q* = 0.01) in domesticated salmon when fed VO diet. The regulator of fatty acid metabolism *pparg-b* was consistently higher expressed in domesticated compared to wild salmon under all diets, but only significantly different for salmon fed FO diet (Figure 5 A).

**Figure 5.**
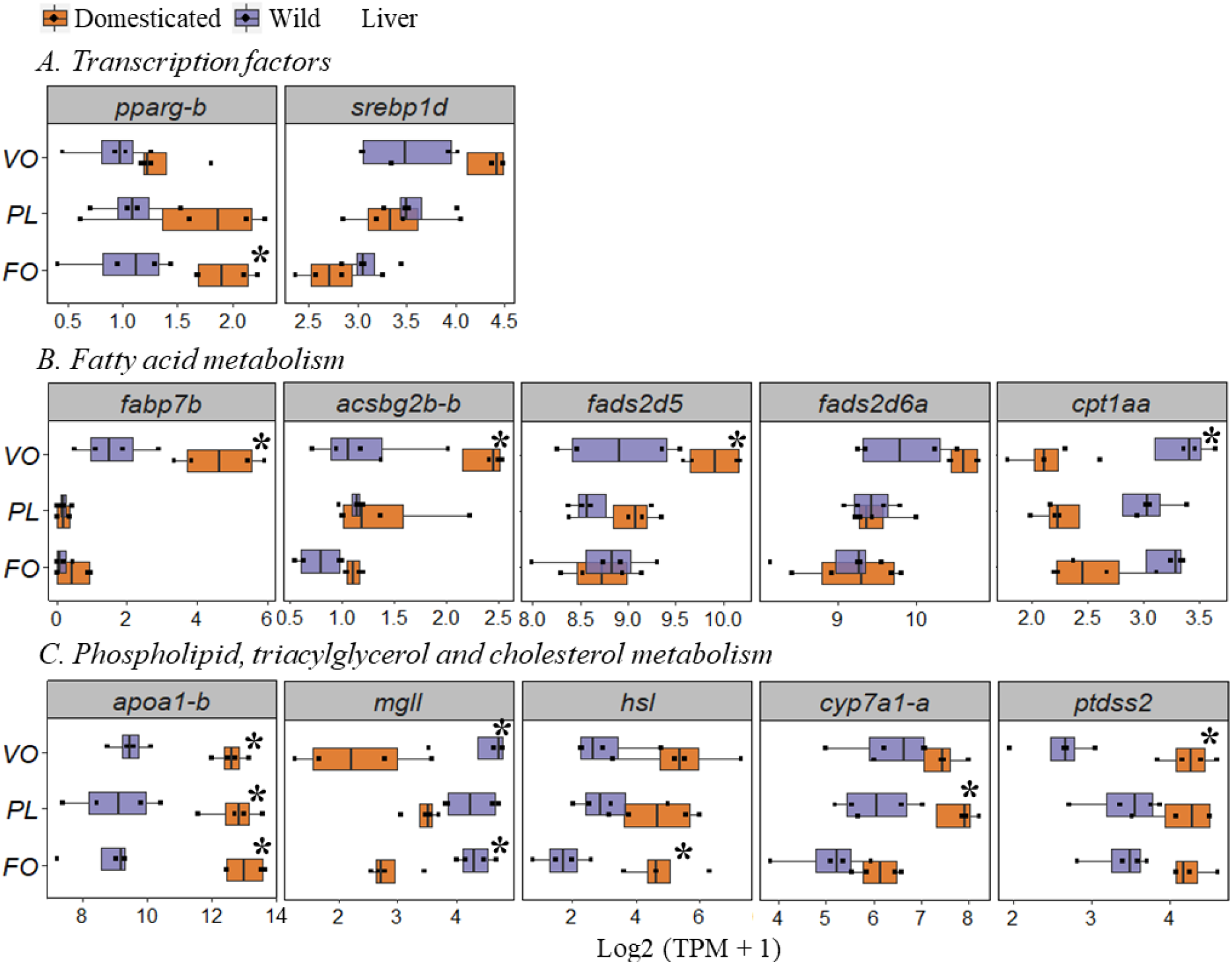
Expression of 14 genes involved in lipid metabolism in liver of wild and domesticated salmon at day 94 after feeding either fish oil (FO), vegetable oil (VO) or phospholipid (PL) diets. Gene expression was shown as Log2 transcript per million plus one (TPM + 1) which was normalized by library size and mRNA length. Asterisk indicates significant (q<0.05 & |Log2FC|>1) different of gene expression between domesticated and wild salmon under each dietary treatment separately.

In addition to the DEGs of fatty acid metabolism, 5 DEGs involved in phospholipid, cholesterol and triacylglycerol metabolism were found between domesticated and wild salmon (Figure 5 C). This included the *apoa1-b* gene involved in lipoprotein synthesis and lipid transport, which was strongly higher expressed in domesticated salmon than wild, regardless of dietary treatment (Log2 FC > 3 & *q*<0.001, see Figure 5 C and Supplementary Table 5). A key gene involved in synthesis of bile acid (*cyp7a1-a*), which is responsible for removal of cholesterol in liver, was higher expressed in domesticated salmon when given PL diet. Gene *ptdss2* involved in synthesis of phosphatidylserine, which is a major phospholipid in salmon, was higher expressed in domesticated salmon than wild fed VO diet (Log2 FC = 1.9, *q* = 0.0008), though similar trend was also found when the fish was given FO (Log2 FC = 1, *q* = 0.09) or PL diet (Log2 FC = 0.9, *q* = 0.2). The expression of *hsl* gene involved in hydrolysing triacylglycerol (stored fat) to diacylglycerol, and diacylglycerol to monoacylglycerol was generally higher expressed in domesticated salmon than wild. On the other hand, the expression of *mgll* involved in hydrolysing monoacylglycerol into free fatty acids was lower expressed in domesticated salmon. (Figure 5 C). In conclusion, the direct comparison of the transcriptomes of domesticated and wild salmon suggests that domestic salmon have boosted expression of genes involved in many aspects of lipid metabolism such as transport, endogenous synthesis and conversion of lipids and fatty acids in both gut and liver (Supplementary table 5).

To further investigate differences in the plasticity of fatty acid metabolism between domesticated and wild salmon, we analysed differences in putative compensatory shifts in gene regulation under diets with low (VO) vs high (FO) levels of LC-PUFA for wild and domesticated salmon separately. These analyses identified 38 DEGs in domesticated and 2 DEGs in wild salmon (Supplementary Table 5). However, only DEGs in domestic salmon (9 genes) were linked to lipid metabolism, specifically involved in fatty acyl-CoA synthesis (2 genes), LC-PUFA synthesis (2 genes), lipogenesis (2 genes), and transcriptional regulation of lipid metabolism (2 genes) (Table 2).

**Table 2.**
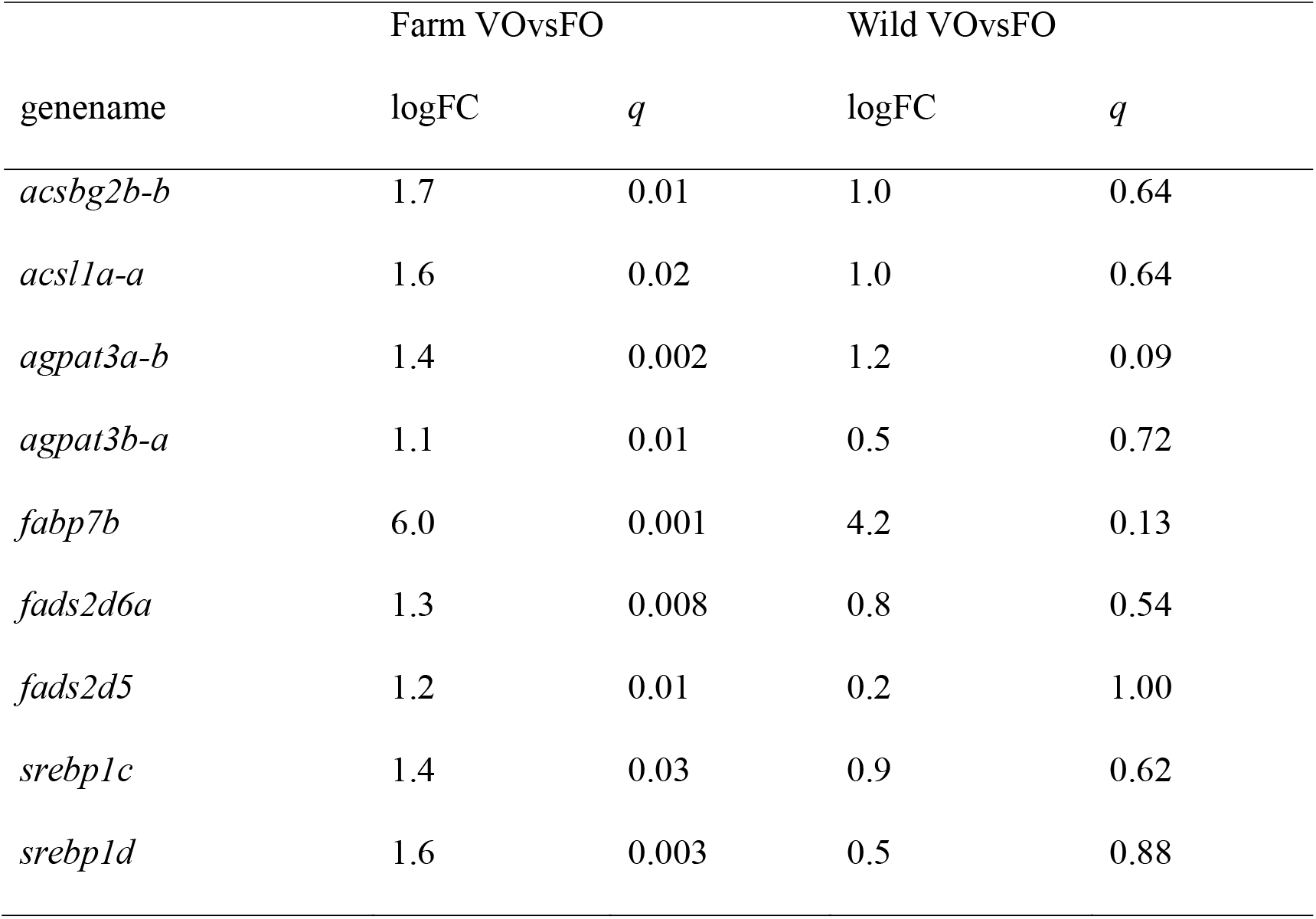
Log2 fold change and adjusted *p* value (*q*) of lipid gene expression in liver of domesticated/wild salmon feeding vegetable oil (VO) diet compared to fish oil (FO).

### 3.5 Comparison of fatty acid composition between domesticated and wild salmon

To assess the differences between fish at the metabolite levels, we also measured fatty acid content in liver and pyloric caeca of wild and domesticated salmon at day 94 (Supplementary Table 6). The results showed that variation in fatty acids composition was generally more driven by diet than strain. About 85% of the fatty acid content in liver and pyloric caeca differed between diets, but only 32% of the fatty acids differed in levels between wild and domesticated salmon (*p*<0.05, Supplementary Table 6). Both wild and domesticated salmon given the VO diet showed higher levels of 18:3n-3 and 18:2n-6 contents in both liver and pyloric caeca but lower contents of the longer chain fatty acids (ARA, EPA & DHA) compared to both FO and PL diets (Figure 6). This pattern was consistent for all three lipid classes analysed (PtdCho, PtdEtn, and TAG). Although the differences in fatty acids content were generally small between wild and domesticated salmon fed the same diet, wild fish contained higher content of 18:2n6 (9.1% in wild *versus* 7.3% in domesticated fish, *p* = 0.06) and 18:3n3 (2.3% *versus* 1.5%, *p* = 0.006) in PtdEtn of liver when fed VO diet. Wild salmon also had higher content of 18:3n3 (2.1% *versus* 1.8%, *p* = 0.04) in PtdCho of liver when fed VO diet. On the other hand, wild salmon had significantly lower content of ARA in both PtdCho (1.6 % *versus* 2.1%, *p* = 0.02) and PtdEtn (3.1% *versus* 4.3%, *p* = 0.02) of liver than wild fish when fed VO diet. No significant differences in DHA and EPA contents were found between domesticated and wild salmon fed the same diets.

**Figure 6.**
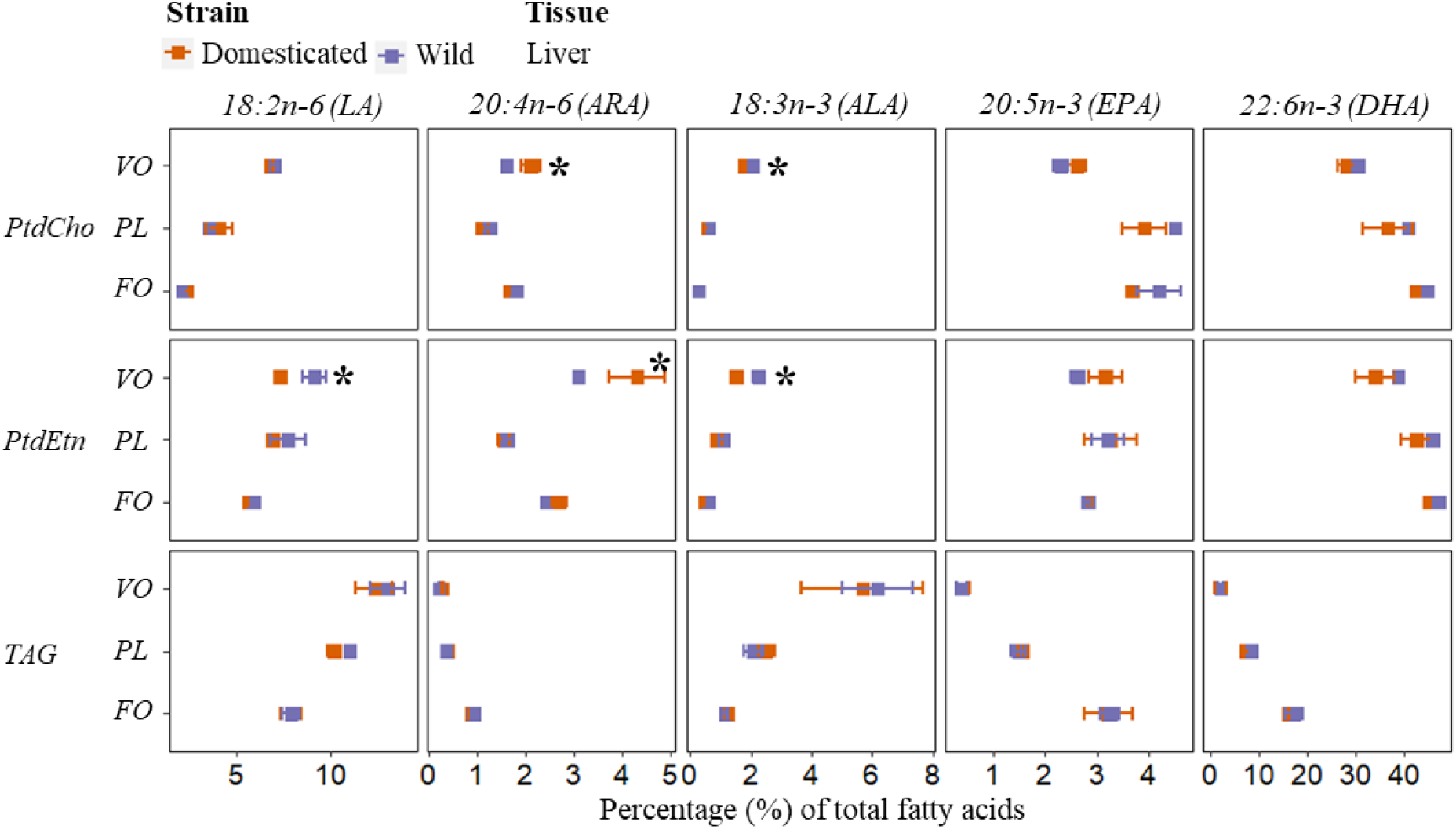
Percentage of liver fatty acid composition in triacylglycerol (TAG), phosphatidylcholine (PtdCho) and phosphatidylethanolamine (PtdEtn) of wild and domesticated salmon fed either fish oil (FO), vegetable oil (VO) and phospholipid (PL) rich diets at day 94. A two-way ANOVA was applied to test the fatty acid differences between fish strains and dietary treatment (strain*diet) separately in each lipid class. Tukey’s HSD post-hoc test was then applied to test the fatty acid difference between each group. Asterix in the figure indicate significant (*p*<0.05) different of fatty acid between domesticated and wild salmon at certain day and dietary treatment. The composition of other fatty acids and their ANOVA test were shown in Supplementary 7.

## 4 Discussion

The present study aimed to improve our understanding of the differences in metabolism of domesticated and wild salmon, with an emphasis on lipid metabolism. We approached this question using comparative analyses of growth, liver and gut transcriptomes, and lipid composition between domesticated and non-domesticated wild salmon given diets with three different lipid composition.

### 4.1 The domestic metabolic syndrome and the link to the circadian clock pathway

As expected, domesticated salmon grew faster, both before and after start feeding, likely reflecting a higher basal metabolic rate (Bicskei, Bron, Glover, & Taggart, 2014) and higher feed intake and feed conversion efficiency (Thodesen et al., 1999) in domestic salmon. In line with this we found a clear difference in expression of genes involved in basal metabolism in liver between domesticated and wild salmon (Figure 3 A). In pyloric caeca, gene expression between wild and domesticated fish was associated with several signaling pathways and glycolipid synthesis (glycosphingolipids and glycophospholipid) which are main components of cell membranes. The promoters of these genes were enriched in TFBS motifs (ETV2, ATOH1, GR GATA-1) known to be involved in intestinal development and cell differentiation (Jedlicka & Gutierrez-Hartmann, 2008; Kanki et al., 2017; Lebenthal & Lebenthal, 1999) (Figure 3 B). One likely explanation for these transcriptomic differences in gut tissue is the difference in growth rates and/or feed intake between domestic and wild fish domesticated (Thodesen et al., 1999).

The enrichment of CLOCK and BHLHB2 motifs in promoters of DEGs (Figure 3) suggests a link between the circadian clock pathway and gene expression differences observed between wild and domesticated salmon (Dunlap, 1999). This finding is interesting as many studies on mammals suggest a functional link between the circadian clock and regulation of feed intake, metabolic rates, and glucose and lipid metabolism (Paschos, 2015; Rudic et al., 2004). The top regulators of circadian oscillations are thought to be CLOCK and BMAL1 transcription factors which form a heterodimer and activate expression of downstream target (Lowrey & Takahashi, 2000). CLOCK/BMAL are involved in regulating (directly or indirectly) a multitude of downstream processes, including genes of metabolism as well as genes that maintain the circadian oscillation of CLOCK/BMAL through negative feedback loops (Lowrey & Takahashi, 2000; Preitner et al., 2002; Takahashi, 2015). Our study does unfortunately not include samples that can shed light on differences between wild and domestic salmon circadian oscillations. However, we find that domesticated salmon generally has lower expression of genes encoding negative regulators of the CLOCK/BMAL (Figure 3 C) (Lowrey & Takahashi, 2000; Preitner et al., 2002; Takahashi, 2015). In this context it is also interesting to note that the PPAR-RXR heterodimer, a major regulator of glucose (Jones et al., 2005) and lipid (Kliewer et al., 1997) homeostasis, is known to be under circadian rhythmicity in salmon (Betancor et al., 2014). The *pparg* gene is consistently higher expressed in domesticated compared to wild salmon (Figure 5) and both PPAR and RXR are predicted to be enriched in promoters of genes differentially expressed in wild and domestic salmon (Figure 3 B). This result is in agreement with the tendency of higher clock gene expression in domesticated fish. It is important to note that all samples used for the gene expression were sampled between morning and noon within a 2h time period. Moreover, fish were raised under constant light and continuous feeding which is known to abolish daily rhythmicity for both *nr1d1* (Betancor et al., 2014) and *cry-2* (Huang, Ruoff, & Fjelldal, 2010). We are thus confident that sampling bias related to daily rhythms has not impacted our results.

In conclusion, we speculate that the ‘domestic metabolic syndrome’ in salmon, characterized by increased basal metabolism and high growth (Bicskei et al., 2014; Tymchuk, Sakhrani, & Devlin, 2009), is partially caused by novel regulation or function of the circadian clock pathway. An interesting hypothesis is that strong selection on ‘fast growers’ during salmon domestication have selected for salmon genotypes that have impacted the circadian clock regulation and thereby induced changes in appetite and metabolism.

### 4.2 Differences in regulation of lipid metabolism between domesticated and wild salmon

In the present study we fed the fish three diets with different fatty acid composition to explore the idea that domestication has changed regulation of lipid metabolism in salmon. We have demonstrated that growth of domesticated salmon was virtually unaffected (Figure 1) by dietary fatty acid composition in the diet, while wild salmon was. This suggest that domesticated salmon have more effective lipid absorption, lipid transport, and better ability to conduct endogenous conversion and synthesis of lipids to compensate for dietary shortage of essential fatty acids. In line with this idea, our study found that *apoa1_2* gene was strikingly higher expressed in liver in domesticated compared to wild salmon, regardless of dietary lipid composition (Figure 5). Although the existence of lipoproteins and their apolipoprotein composition is not well described in fish, it is suggested that they should be similar to mammals (Jin et al., 2018). The higher expression levels of *apoa1_2* transcripts could therefore be one factor that contributes to fast growth of domesticated salmon, as this gene is key to the lipoprotein assembly and thereby transport of lipids between liver and other tissues such as muscle and adipose tissue (Xu et al., 2013). We also observed higher expression of hormone sensitive lipase gene (*hsl*) in liver of domesticated salmon than in wild. This suggests that domesticated salmon has higher ability of hydrolysing triacylglycerol, diacylglycerol and cholesterol ester into monoacylglycerol and free fatty acids (Kraemer & Shen, 2002; Quiroga & Lehner, 2012). The hydrolysed monoacylglycerol and fatty acids were more likely to be used in re-synthesis of lipids rather than further degraded to produce energy, because key genes involved in fatty acid degradation, the *mgll* gene involved in hydrolysing monoacylglycerol and *cpt* genes involved in transporting fatty acids into the mitochondria for β-oxidation, were lower expressed in domesticated salmon than wild.

Wild fish fed the PL diet had the fastest growth rate, which supports previous findings that dietary inclusion of PL provides juvenile salmon better growth and development (Poston, 1990; Taylor et al., 2015). This is likely a consequence of PL promoting the absorption and transport of dietary lipid especially LC-PUFA (Carmona-Antonanzas, Taylor, Martinez-Rubio, & Tocher, 2015; Rolf Erik Olsen, Tore Dragnes, Myklebust, & Ringø, 2003; Y. Olsen et al., 2014a; D. R. Tocher, E. Å. Bendiksen, P. J. Campbell, & J. G. Bell, 2008). Young salmon have low capacity of *de-novo* synthesis of PL and may therefore struggle to maintain sufficient lipid levels for high growth (Carmona-Antonanzas et al., 2015; Jin et al., 2018; Rolf Erik Olsen et al., 2003; Y. Olsen et al., 2014b; D. R. Tocher, E. A. Bendiksen, P. J. Campbell, & J. G. Bell, 2008). In the present study, we identified higher expression of 2 phosphate cytidyltransferase genes (*pcyt2c-a* and *pcyt2c-b*) in pyloric caeca of domesticated than wild salmon only when fish was given FO or VO diets, while no difference of gene expression was found when the fish was fed the PL diet. This suggests that domesticated salmon can increase the expression of genes involved in PL biosynthesis to compensate for the dietary shortage of PL, while such ability is limited in wild salmon at the same age. The expression of phosphatidylserine synthase gene (*ptdss2*) was generally higher expressed in both pyloric caeca and liver of domesticated salmon regardless of dietary treatment. The reason for the higher requirements of phosphatidylserine in domesticated salmon is not clear. One of the reasons could be that phosphatidylserine is a constituent of the cell membrane, which is required when there is a potential higher level of cell proliferation in domesticated salmon.

Domesticated salmon are more adapted to the VO diet compared to wild strains. This was clearly revealed in the growth data where dietary VO significantly affected growth and final weight of wild salmon, but had no effect on the growth of domesticated salmon. It is generally assumed that a VO enriched diet can induce the expression of LC-PUFA synthesis genes in salmon (Stubhaug et al., 2005; Zheng et al., 2004). However, the key genes in the LC-PUFA synthesis pathway, *fads2d5* and *fads2d6a* were up-regulated only in domesticated salmon when fed the VO diet. This suggests that domesticated salmon is able to modify its fatty acid metabolism to compensate for the shortage of essential LC-PUFA in the diet, while such ability is very low in wild salmon. Although previous generations of domesticated salmon were fed VO diet only after transition to seawater, we still observed shifted expression of lipid metabolism domesticated but not wild salmon at freshwater stage. This suggests that the adaption to VO diet in domesticated salmon is independent of life stages. The present study showed higher contents of DHA, ARA and EPA and lower contents of 18:3n3 and 18:2n6 in phospholipid of liver in domesticated salmon than wild only when they were given VO diet. This was most likely an effect of higher LC-PUFA synthesis ability in domesticated salmon. Sterol regulatory binding protein 1 (SREBP1) is believed to be the key regulator of fatty acid metabolism and their expression is influenced by LC-PUFA level in the cell (Datsomor et al., 2019). We observed an up-regulation of the *srebp1d* gene in domesticated salmon, but not in wild fish when fed VO diet. Acyl-CoA synthase (*acsbg2b-b*) genes were also up-regulated, which was believed to be targeted by SREBP1 transcription factor (Datsomor et al., 2019).

In conclusion, the present study has provided a broad overview on transcriptomic and fatty acids differences in wild and domesticated salmon fed either FO, VO or PL enriched diets during early stages of development. The higher growth and development of domesticated salmon was in agreement with a combination of various genetic advantages including better uptake, transport and endogenesis of lipids. This was associated with altered circadian clock regulation between domesticated and wild salmon, which could possibly explain the changes in appetite and metabolism. Moreover, domesticated salmon had higher plasticity on gene expression when fed a VO diet with less essential LC-PUFA, while this ability is very low in wild salmon as the growth of fish significantly dropped for the VO diet. However, further experiment on circadian oscillations is required to support the differential expression of circadian genes found in present study. Other layers of genetic and biochemical information are also needed to get an in-depth and complete understanding of divergence of salmon after domestication. We suggest that future studies should focus on comparing metabolites, lipidome and proteome between domesticated and wild salmon.

## Acknowledgements

The design and running of the experiment was supported by non-specific grants from Department of Biology, Norwegian University of Science and Technology (NTNU). The domesticated salmon eggs were kindly provided by AquaGen AS with assistance from Dr. Maren Mommens. The RNA-Seq and data analysis were financed by the Research Council of Norway (GenoSysFat, grant number 244164) and (DigiSal, grant number 248792). The sequencing service was provided by the Norwegian Sequencing Centre, a national technology platform hosted by the University of Oslo. The herring roe used in PL diet was kindly provided by Erik Løvaas from Marine BioExploitation AS. We also would like to thank Jostein Ervik for rearing the fish and Eleni Nikouli for the help on sampling. Thank you Torfinn Sparstad, Signe Dille Løvmo, Hanne Hellerud Hansen and Centre for Integrative Genetics (CIGENE) for RNA-Seq sample preparation. Thank you Dr. Gareth Gillard for pre-processing the RNAseq data, mapping reads to salmon genome and acquire count reads. We also thank the China Scholarship Council for providing financial support to Yang Jin for his PhD study.

## Data Accessibility Statement

Raw sequences are publicly available on ArrayExpress under accession number E-MATB-8306.

## Author Contributions

Yang Jin, Yngvar Olsen, Rolf Erik Olsen, Simen Rød Sandve and Olav Vadstein designed and performed the research. Yang Jin and Thomas Nelson Harvey performed the transcriptomic analysis. Yang Jin and Keshuai Li performed the lipid and fatty acid analysis. Jon Olav Vik and Simen Rød Sandve guided the transcriptomic analysis and revised the manuscript. Yngvar Olsen and Rolf Erik Olsen guided the lipid analysis and revised the manuscript. Mari-Ann Østensen and Nina Santi provided input on the experimental design, carried out the experiment and sampling and reviewed the manuscript. All authors participated in the revision of this paper by providing comments and editing.

