## Supplementary Figure 1 for "Comparative transcriptomics reveals domestication-associated features of Atlantic salmon lipid metabolism"

**Expression of circadian genes between farmed and wild salmon**

1. **Liver (day 65 and 94 after initial feeding)**

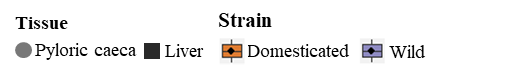

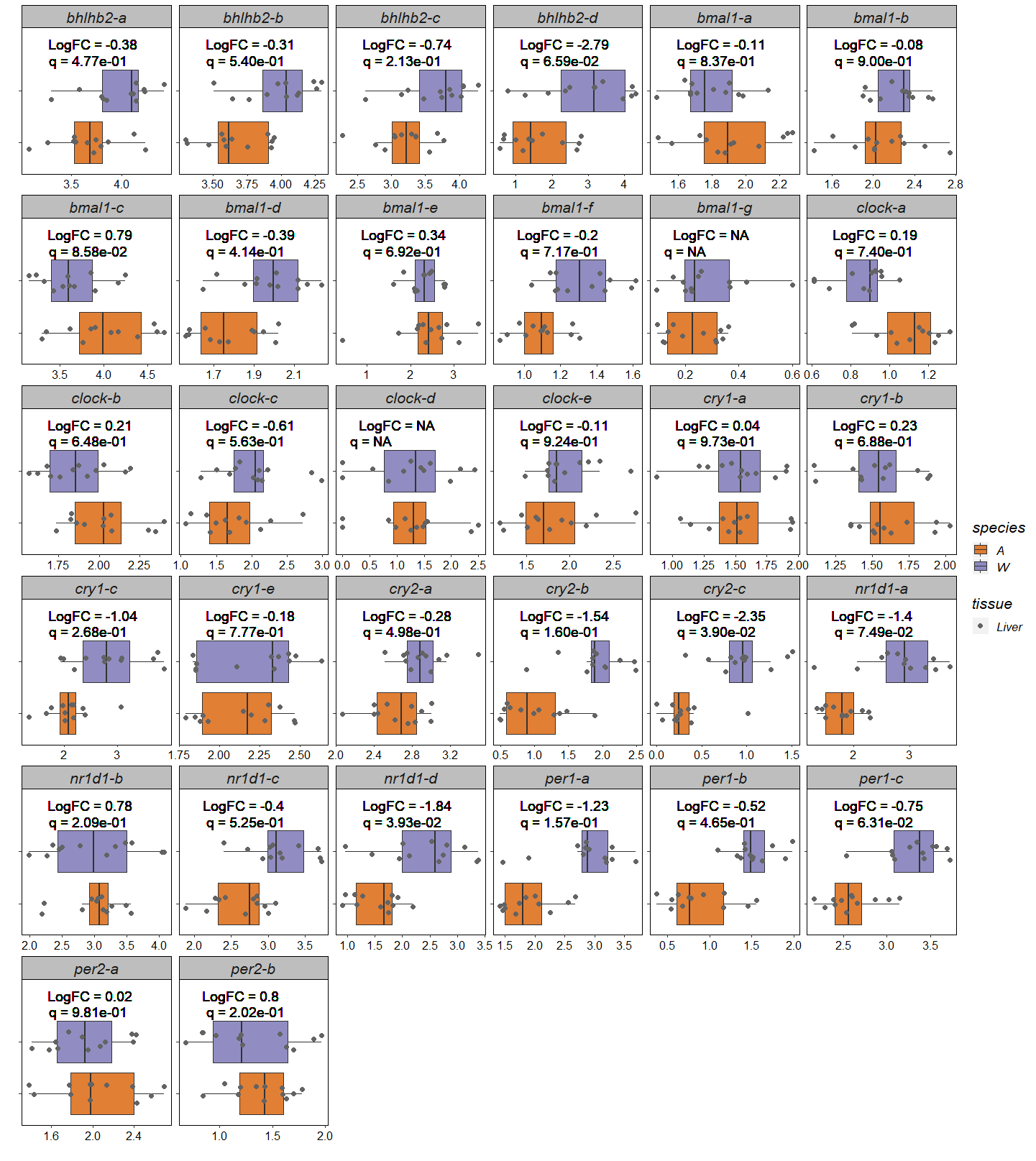

2. **Pyloric caeca (day 94 after initial feeding)**

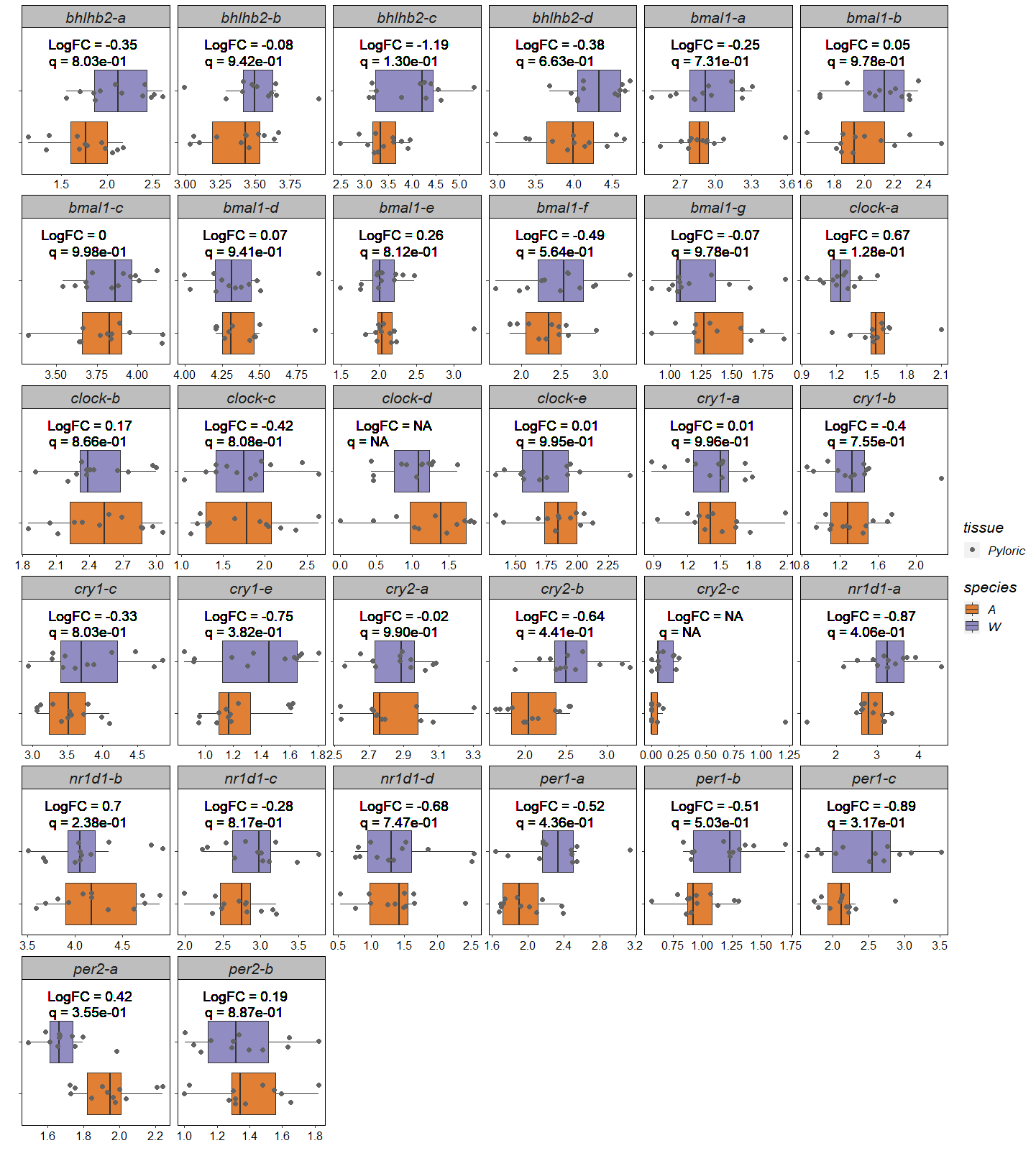

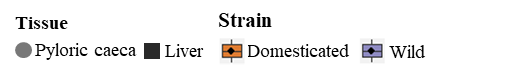
